# Multiscale heterogeneity of white matter morphometry in psychiatric disorders

**DOI:** 10.1101/2024.08.04.606523

**Authors:** Ashlea Segal, Robert E Smith, Sidhant Chopra, Stuart Oldham, Linden Parkes, Kevin Aquino, Seyed Mostafa Kia, Thomas Wolfers, Barbara Franke, Martine Hoogman, Christian F. Beckmann, Lars T. Westlye, Ole A. Andreassen, Andrew Zalesky, Ben J. Harrison, Christopher G. Davey, Carles Soriano-Mas, Narcís Cardoner, Jeggan Tiego, Murat Yücel, Leah Braganza, Chao Suo, Michael Berk, Sue Cotton, Mark A. Bellgrove, Andre F. Marquand, Alex Fornito

## Abstract

**Background:** Inter-individual variability in neurobiological and clinical characteristics in mental illness is often overlooked by classical group-mean case-control studies. Studies using normative modelling to infer person-specific deviations of grey matter volume have indicated that group means are not representative of most individuals. The extent to which this variability is present in white matter morphometry, which is integral to brain function, remains unclear.

**Methods:** We applied Warped Bayesian Linear Regression normative models to T1- weighted magnetic resonance imaging data and mapped inter-individual variability in person-specific white matter volume deviations in 1,294 cases (58% male) diagnosed with one of six disorders (attention-deficit/hyperactivity, autism, bipolar, major depressive, obsessive–compulsive and schizophrenia) and 1,465 matched controls (54% male) recruited across 25 scan sites. We developed a framework to characterize deviation heterogeneity at multiple spatial scales, from individual voxels, through inter-regional connections, specific brain regions, and spatially extended brain networks.

**Results:** The specific locations of white matter volume deviations were highly heterogeneous across participants, affecting the same voxel in fewer than 8% of individuals with the same diagnosis. For autism and schizophrenia, negative deviations (i.e., areas where volume is lower than normative expectations) aggregated into common tracts, regions and large-scale networks in up to 35% of individuals.

**Conclusions:** The prevalence of white matter volume deviations was lower than previously observed in grey matter, and the specific location of these deviations was highly heterogeneous when considering voxel-wise spatial resolution. Evidence of aggregation within common pathways and networks was apparent in schizophrenia and autism but not other disorders.

## Introduction

The brain’s white matter comprises densely packed myelinated axons that inter connect distinct regions of grey matter and play a critical role in healthy brain function (1,2). Damage to white matter can disrupt coordinated neuronal activity, leading to clinical symptoms of mental illness. Accordingly, magnetic resonance imaging (MRI) studies report diverse white matter abnormalities in different psychiatric diagnoses (3–10).

These abnormalities are commonly revealed by traditional statistical approaches that rely on comparisons of group means, which can overlook the tremendous clinical and biological inter-individual variability that characterizes psychiatric illness (11–14). Normative modelling (15,16) offers a framework for characterizing this variability by allowing one to quantify the extent to which individuals deviate on a given phenotype from normative expectations given relevant demographic characteristics, such as age and sex (15–17). This approach has been applied to a range of MRI-derived phenotypes in different neurodegenerative and psychiatric disorders (18–26).

However few studies have used normative modelling to investigate the heterogeneity of white matter deviations in psychiatric disorders but they have generally agreed that no more than 20% of individuals with the same diagnosis show a significant WMV deviation in the same white matter location or tract (20,23,24,26). While this heterogeneity aligns with the noted clinical variability of psychiatric populations (11), it raises an important question: how can individuals be assigned the same diagnosis if their brain deviations are so heterogeneous?

One possibility is that deviations occurring in different anatomical locations but aggregate within common, spatially distributed brain networks. Our recent work on grey matter volume (GMV) showed that while individuals with attention-deficit hyperactivity disorder (ADHD), autism spectrum disorder (ASD), bipolar disorder (BP), major depressive disorder (MDD), obsessive-compulsive disorder (OCD), or schizophrenia (SCZ) had minimal overlap in GMV deviation locations, these deviations were often within shared functional networks (22). We hypothesized that a similar pattern might occur in white matter tracts; i.e. WMV deviations may not necessarily co-localize in space with the same diagnosis, but they may nonetheless implicate a common white matter tract, brain region, or extended brain network.

To test this hypothesis, we quantified the anatomical heterogeneity of WMV deviations in 1465 healthy controls and 1294 cases diagnosed with one of six psychiatric conditions (ADHD, ASD, BP, MDD, OCD, SCZ) (22). We developed a framework to characterize WMV deviations at multiple scales of analysis to determine the degree to which spatially heterogeneous deviations may accumulate within common white matter pathways, connections tied to individual brain regions, or spatially extended brain networks.

## Materials and methods

The study was approved by the local ethics committee of the site contributing each dataset, and written informed consent was obtained from each participant. The overall study was approved by the Monash University Research Ethics Committee (Project ID: 23534). Data processing was conducted using Multi-modal Australian ScienceS Imaging and Visualisation Environment (MASSIVE) (27).

### Participants

A final sample of 1294 cases and 1465 controls was obtained from a larger cohort drawn from 14 separate, independently acquired datasets and 25 scan sites following various exclusion criteria (see *Supplementary Material: Participants*) (22). All cases were diagnosed according to clinical judgement and diagnostic instruments as per individual study procedures. All individuals were aged between 18-64 years old. Basic demographic details of the final sample are provided in Table 1. For a comprehensive overview inclusion criteria and demographic details for each site see relevant references in Table 1 and *Supplementary Material: Data Availability*.

**Table 1.**
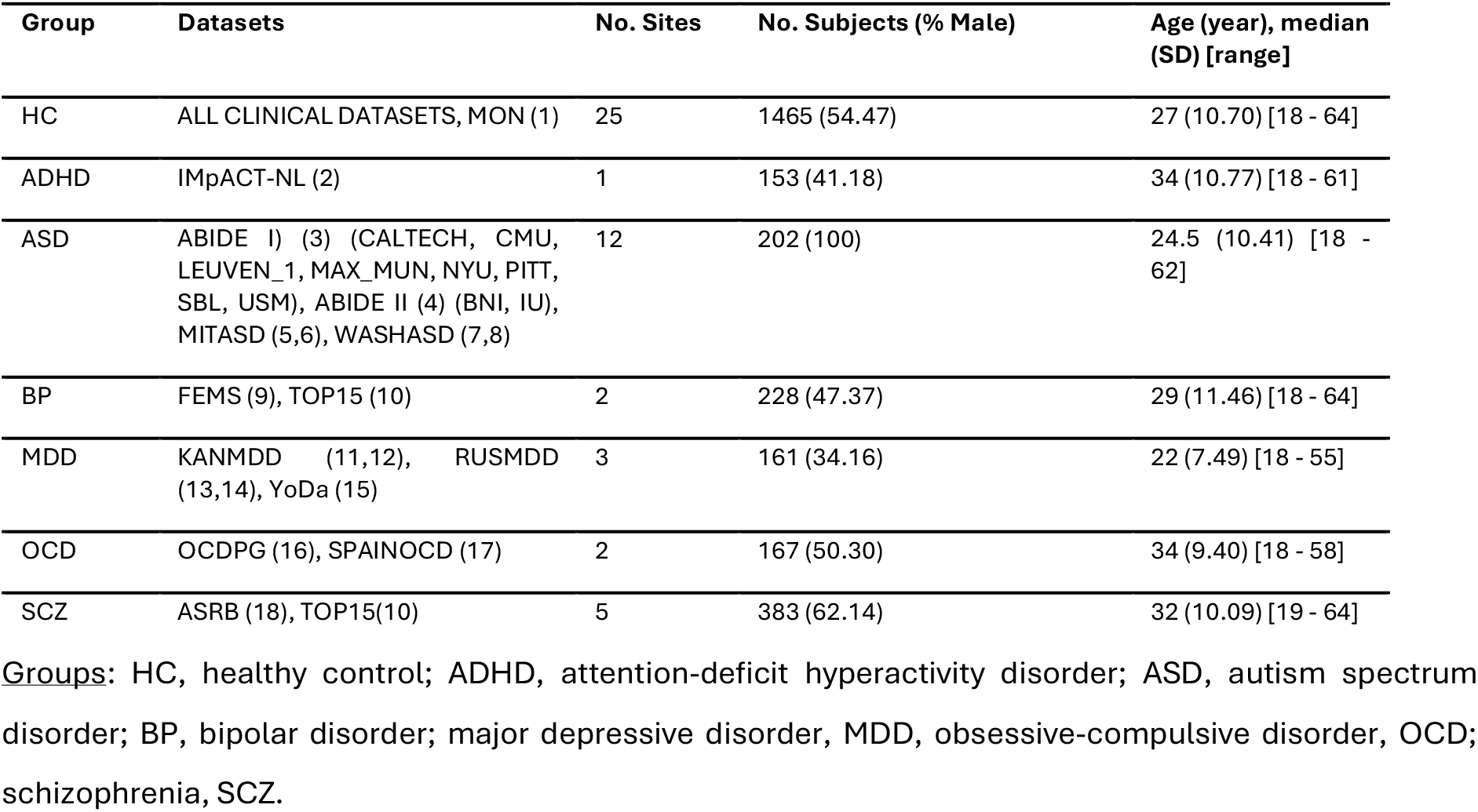
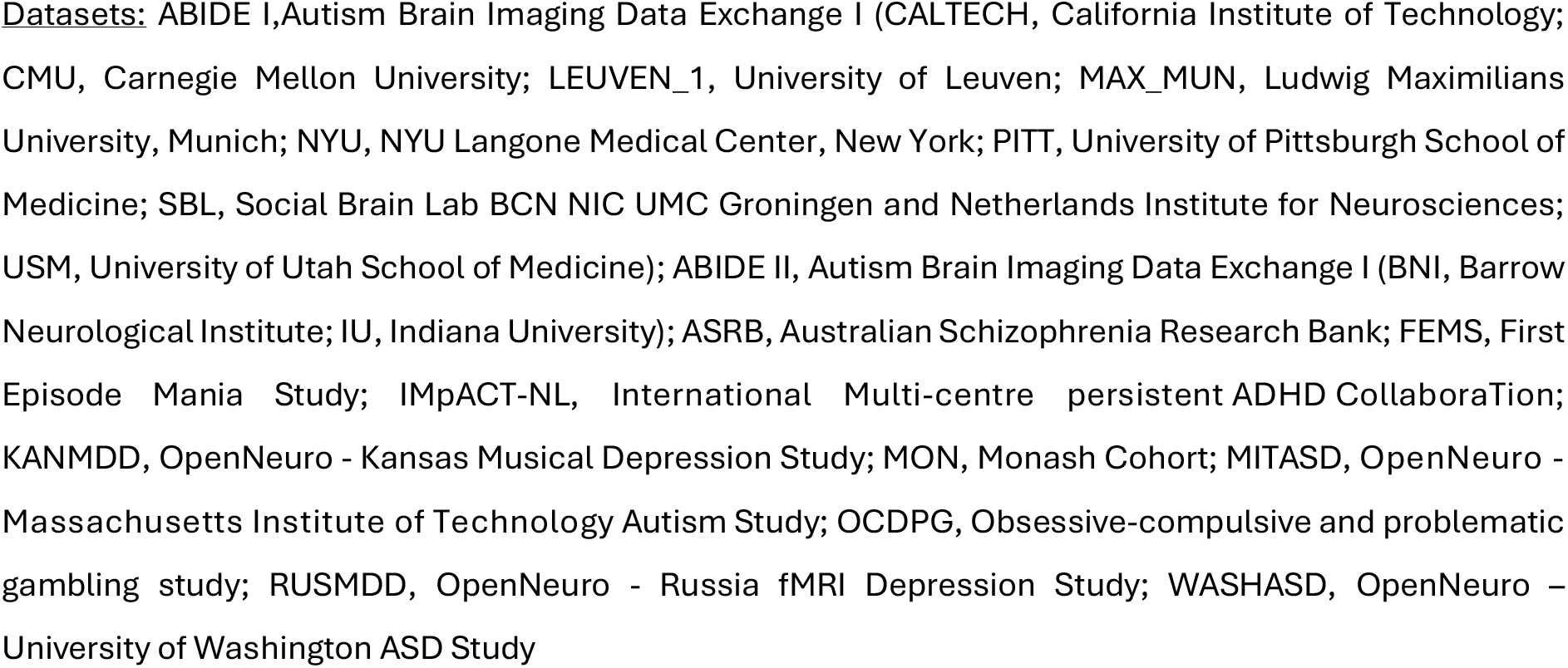
Summary of demographic data of the clinical and control groups

### Analysis Overview

We characterized neural heterogeneity across various “scales” of network resolutions: individual white matter volume (WMV) voxels, white matter tracts (i.e., connections between pairs of grey matter regions), brain regions, and canonical functional networks. To map WMV heterogeneity at the voxel scale, we used normative models to create person-specific voxel-wise deviation maps, showing how each person’s WMV deviates from normative predictions (Figure 1a-e). This involved (1) estimating voxel-wise WMV for each individual, (2) quantifying deviations from normative expectations, and (3) applying thresholds to identify deviant WMV clusters.

**Figure 1.**
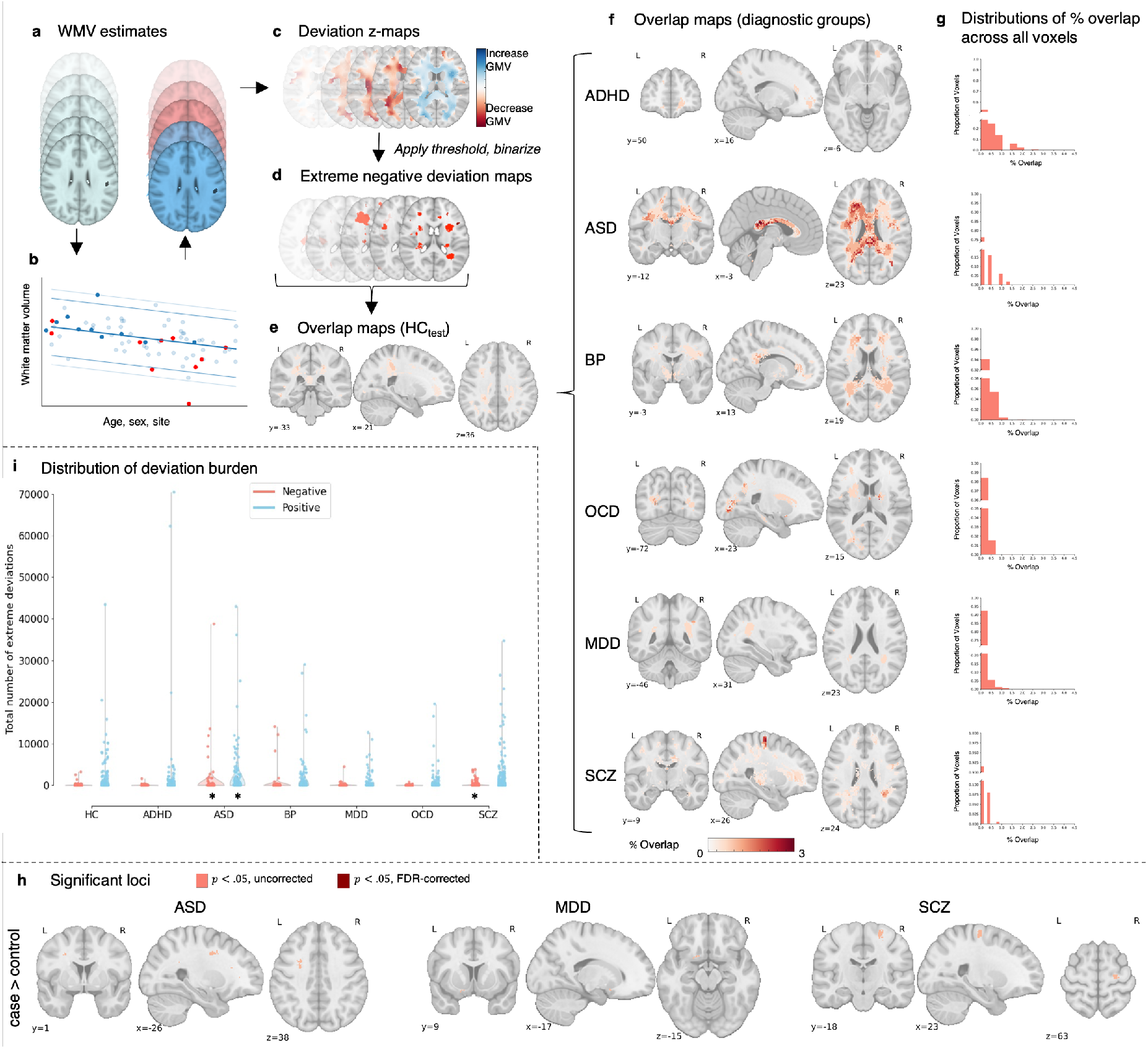
Voxel-wise heterogeneity of extreme WMV deviations. (**a-g**) *Workflow to characterise voxel-scale WMV heterogeneity*: (**a)** WMV maps of individuals in the HCtrain cohort were used to train a normative model to make voxel-wise WMV predictions given someone’s age, sex, and scan site. The predictions for the held-out control (HCtest; red) and clinical (cases; dark blue) participants were then compared to the empirical WMV estimates. (**b**) Shows an example of model predictions for a given voxel, with light blue dots representing individuals in the training set, dark blue representing held-out controls (HCtest), and red representing participants in the clinical group. (**c**) For each individual, deviations from model predictions are quantified as z- scores and scaled by model uncertainty to yield a deviation z-map. (**d**) This deviation map is then thresholded at z <- 2.6 to identify regions showing extreme negative deviations. (**e**) For the HCtest and each clinical group, we quantified the proportion of individuals showing an extreme deviation in a given voxel, yielding an extreme deviation overlap map. (**f-g**) *Spatial overlap of extreme negative WMV deviations:* (**f**) Spatial maps showing the extreme deviation overlap map for each clinical group. (**g**) Histograms showing the distribution of overlap percentages observed across all voxels. Note that the y-axis is broken to show the distribution of non-zero values. (**h**) Spatial maps showing voxels with significantly greater overlap in cases compared to controls in extreme negative deviations (*p* < .05, two-tailed, cases>controls). There were no extreme WMV deviations with significantly greater overlap in ADHD, OCD, and OCD compared to controls at p<.05 uncorrected, no extreme WMV deviations with significantly greater overlap in ASD, MDD, and SCZ at p<.05 FDR-corrected, and no extreme WMV deviations with significantly greater overlap in controls compared to cases. (**i**) Distribution of positive (Z > 2.6; blue) and negative (Z <-2.6; red) deviation burden scores (i.e., the total number of extreme deviations) in each diagnostic group. *\** Indicates clinical groups showing a statistically significant difference in extreme deviation burden compared to the HCtest group (Mann Whitney U-test, *p* < .05, one- tailed). Steps (**d-h**) were subsequently repeated for positive deviations, with the deviation map is thresholded at z > 2.6 to identify regions showing extreme positive deviations.

Next, we examined whether WMV deviations affected common axonal pathways linking similar brain regions or networks, using an independent normative diffusion tractography dataset. This generated a dysconnectome map for each person, representing affected pairwise connections (Figure 2a-f). We then assessed if these connections were concentrated in specific brain regions or large-scale networks (Figure 3a-e). This approach tested the hypothesis that while disorders show substantial heterogeneity in the anatomical location of WMV deviations, these deviations commonly affect specific axonal pathways linking brain regions or networks.

**Figure 2.**
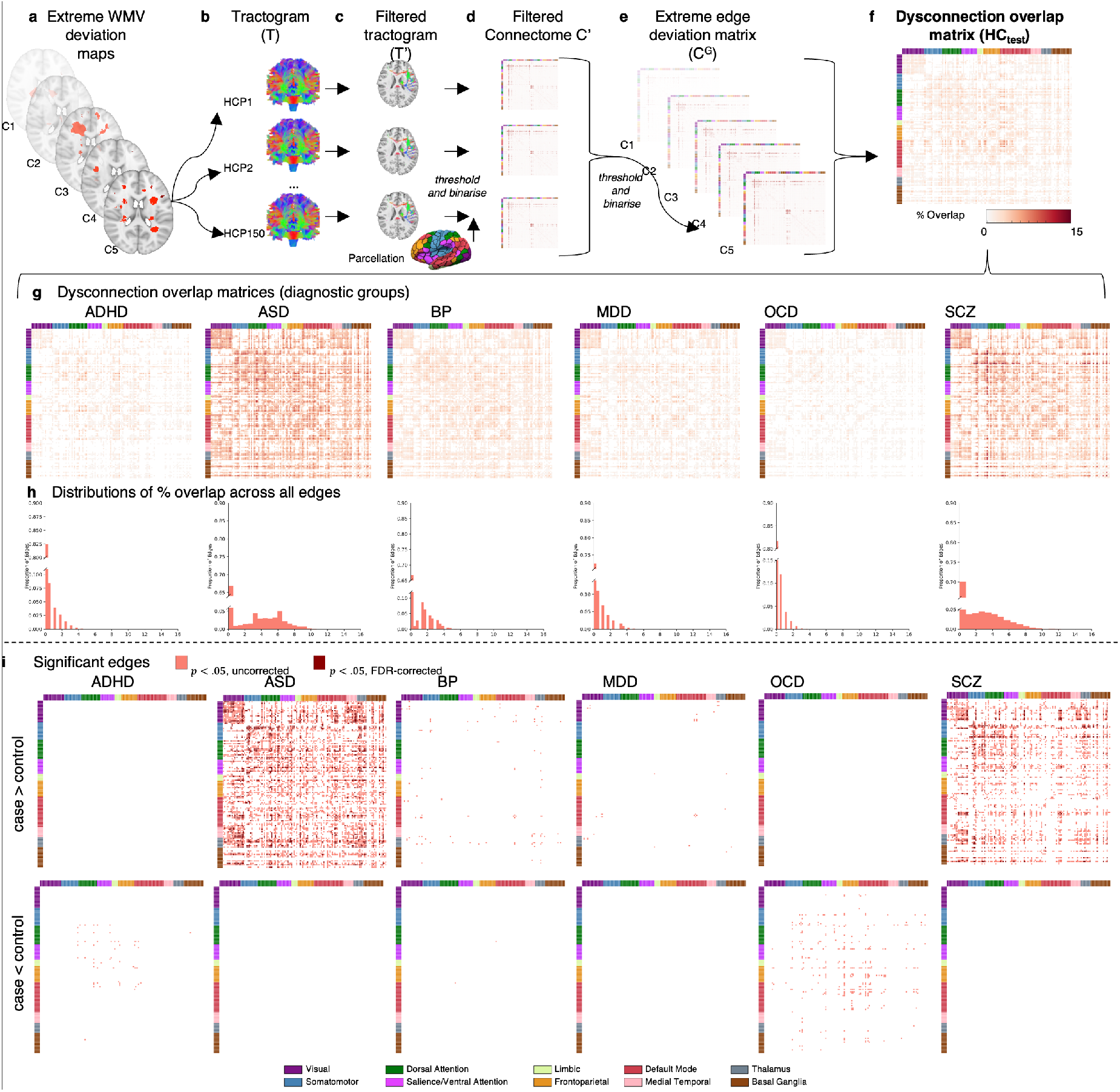
Inter-regional connections affected by extreme negative WMV deviations. (***a-f****) Workflow to characterise pairwise tract-scale WMV heterogeneity*: (**a**) For each participant in HCtest and each clinical group, we used the extreme deviation map to (**b-c**) filter a tractogram obtained for each participant in an independent sample of controls (HCP150). (**d**) This resulted in an adjacency matrix of the pairwise connections affected by a deviation in each case and control, and each person in the HCP150 sample, denoted C’. We thresholded (number streamlines >= 10) and binarised each matrix. (**e**) For each participant in HCtest and each clinical group, we aggregated across the HCP150 adjacency matrices, resulting in a consensus matrix for each case and control, denoted 𝐶^!^. We then thresholded (50% or 75% of HCP150 cohort) and binarised this matrix to identify structural connections affected by WMV extreme deviations for each case and control. (**f**) For the HCtest and each clinical group, we quantified the proportion of individuals showing structural connections affected by WMV extreme deviations in each edge, yielding an edge-scale dysconnection overlap matrix. (**g-i**) *Spatial overlap of edge-scale dysconnection*: (**g**) Matrices showing tract-scale dysconnection for each clinical group (**h**) Histograms showing the distribution of overlap percentages observed across all inter-regional connections. Note that the y-axis is truncated to better illustrate the shape of the distribution. (**i**) Matrices showing edges structurally connected to extreme negative WMV deviations (Z < - 2.6, cluster size threshold=10 voxels) with significantly greater overlap in cases, compared to controls (top), and significantly greater overlap in controls, compared to cases (bottom, *p* < .05, two-tailed). All matrices are sorted according to the Yeo et al(34) network parcellation, followed by the medial temporal lobe, thalamus, and basal ganglia systems.

**Figure 3.**
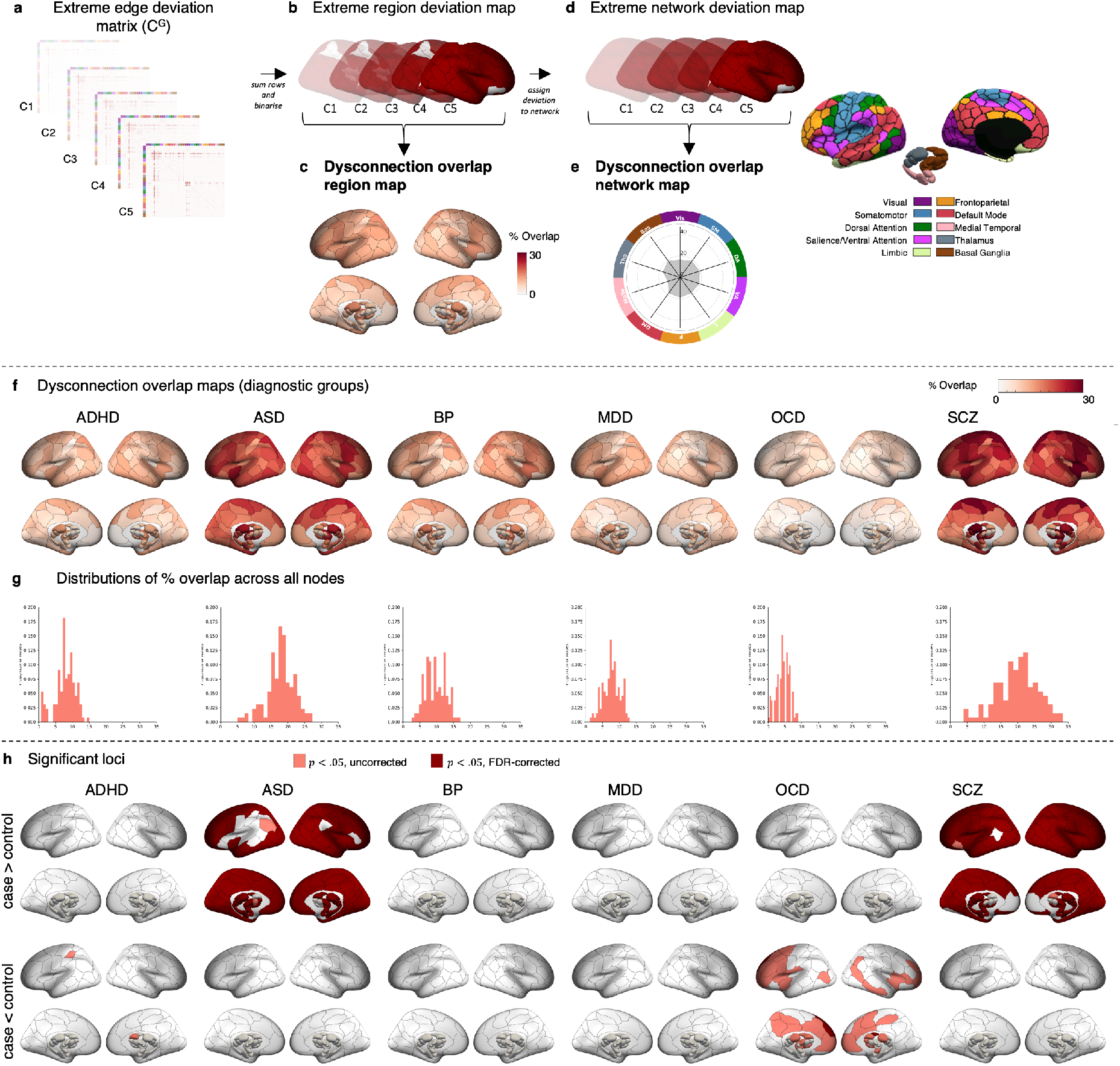
Regions and functional networks attached to connections affected by extreme negative WMV deviations. (***a-e****) Workflow to characterise region- and network-scale WMV heterogeneity. Region scale:* (**a**) For each individual in HCtest and each clinical group, we used the binarised consensus adjacency matrix (**b**) to generate a map of affected grey matter regions. We then quantified the proportion of individuals in each group showing edge-scale dysconnection to each region, yielding a region-scale dysconnection map (**c**). *Network scale:* (**d**) For each individual in the HCtest and each clinical group, we assigned each brain region showing region- scale dysconnection to one of 10 canonical cortical functional networks. An entire network was considered deviant if it contained at least one region linked to an extreme deviation. (**e**) We quantified the proportion of individuals in each group showing a deviation within each network, yielding a network scale dysconnection map. (**f-h**) *Spatial overlap of region-scale dysconnection*: (**f**) Spatial maps quantifying the proportion of individuals showing significant structural connectivity in each region, yielding a region-scale dysconnection map for each diagnostic group. (**g**) Histograms showing the distribution of overlap percentages observed across all regions. (**h**) Statistical maps showing regions structurally connected to extreme negative WMV deviations (z < - 2.6, cluster size threshold=10 voxels) with significantly greater overlap in cases, compared to controls (top), and significantly greater overlap in controls compared to cases (bottom, *p* < .05, two-tailed).

### Voxel-scale WMV heterogeneity

We estimated voxel-wise WMV using voxel-based morphometry (VBM) of T1-weighted anatomical MRI scans (28,29) as per (22) (see *Supplementary Materials: Anatomical data*). Details of the VBM procedure, image acquisition, and quality assurance procedures for the T1-weighted images are detailed in (22).

We used normative modelling to obtain person-specific WMV deviation maps relative to a model, based on Bayesian linear regression (BLR) with likelihood warping (30), of population expectations for voxel-wise WMV variations (see https://github.com/amarquand/PCNtoolkit, version=0.21) (15,17). Following stratification of the sample into training (HCtrain, *n* = 1,196) and test subsets (clinical data, *n* = 1,294; HCtest, *n* = 269) (Figure 1a) (see *Supplementary Materials: Normative modelling* and (22)), we fitted separate models to each white matter voxel to predict WMV as a function of age, sex, and site (Figure 1b) (see *Supplementary Materials: Normative modelling*). We then computed deviation 𝑧*-*maps for each individual (Figure 1c), which quantify the extent to which individual voxel-wise WMV estimates deviate from the model prediction (see (15,16) for details). See *Supplementary Materials: Evaluation of model performance* for the evaluation of model performance. Extreme deviations were defined as clusters of 10 contiguous voxels with deviation scores of |*z*| > 2.6 (i.e., *p* < .005) (22,23) (Figure 1d). See *Supplementary Materials: Normative modelling* for details.

To characterize the inter-individual heterogeneity of deviation locations, we calculated the proportion of individuals showing an extreme deviation within each white matter voxel, separately for positive and negative deviations (Figure 1e-f) (22). We then used permutation testing to test for statistically significant differences in the proportions of cases and controls (HCtest) showing an extreme deviation in each voxel (see *Supplementary materials: Characterizing voxel-wise heterogeneity of extreme deviations for details*).

### Tract-scale WMV heterogeneity

The network context of WMV deviations was evaluated with respect to a normative connectome estimated in an independent healthy unrelated cohort of 150 diffusion-weighted MRI scans (71 males, aged 21-35 years) from the S900 release of the Human Connectome Project (HCP150) (31). The connectome mapped 8646 tracts linking each pair of 100 cortical and 32 subcortical regions, as defined using established atlases (32,33) (see *Supplementary Materials: Diffusion-weighted imaging acquisition parameters and processing* for details).

We produced a 132×132 dysconnection overlap matrix for each group using extreme deviation maps for each participant and tractograms from the HCP150 cohort. Each matrix element represents the fraction of participants whose WMV deviation intersected the tract linking regions i and 𝑗, with separate matrices for positive and negative deviations. Tract-scale heterogeneity was quantified using a permutation-based inferential procedure similar to the analysis of voxel-scale deviations. See Figure 2 and *Supplementary Materials: Streamline region assignments* and *Supplementary Materials: Identifying streamlines intersecting with WMV deviations for details*.

### Region-scale WMV heterogeneity

To determine whether any affected tracts were preferentially linked to specific brain regions, we generated a map of grey matter regions attached to at least one affected tract in the dysconnectome matrix of each individual in the clinical and HCtest groups, (Figure 3a). The resulting person-specific binary regional maps (Figure 3b), obtained separately for positive and negative deviations, were then aggregated across individuals to obtain group-specific overlap maps quantifying, at each region, the proportion of people within a given group for whom at least one affected connection was attached to that region (Figure 3e). We next evaluated case-control differences in overlap proportions for each region using the same procedures as those used in the voxel-wise and tract-scale analyses (see *Supplementary Materials* for details).

### Network-scale WMV heterogeneity

We next examined the degree to which WMV deviations aggregated within distinct functional networks. Using extreme region deviation maps (Figure 3b), we assigned each cortical region to one of seven canonical functional cortical networks using a well-validated network parcellation (34) and assigned each subcortical region to either medial temporal lobe (amygdala and hippocampus), thalamus, or basal ganglia (nucleus accumbens, globus pallidus, putamen, caudate nucleus), as done previously (33), resulting in a total of 10 distinct functional networks (Figure 3d).

For each network, we then estimated the proportion of individuals in each diagnostic group with at least one extreme deviation in any region of each network, creating group-specific dysconnection networks for positive and negative deviations separately (Figure 3e). Case-control differences in overlap proportions for each network were evaluated using the same procedures as those outlined in previous analyses (see *Supplementary Materials* for details). We repeated the analysis using a 20-network parcellation (17 cortical networks and 3 subcortical groups).

## Results

### Normative Modelling

The model evaluation metrics indicated good model fits and successful removal of site effects. We only excluded 1.70% of voxels due to poor model fit (see Table S2 and *Supplementary materials: Model evaluation*).

Figure 1 i shows the distribution of positive and negative person-specific deviation burden scores (i.e., the total number of extreme deviation voxels identified in a person) across individuals, stratified by diagnostic group. In total, 21.56% of controls showed at least one negative extreme deviation; for cases, between 13.17% (OCD) and 35.77% (SCZ) showed at least one negative extreme deviation. For positive deviations, 60.59% of controls showed at least one extreme value, compared to a range of 50.93% (MDD) to 71.78% (ASD) for the clinical group (Table 2). People with ASD and SCZ showed a higher median extreme negative deviation burden compared to held-out control group (HCtest) (nonparametric rank sum test, *p*<.001, one-tailed), and people with ASD also showed a higher median extreme positive deviation burden compared to HCtest (*p*=.001, one-tailed). No other significant case-control differences were identified.

**Table 2.** Deviation burden at the voxel-scale

|  | Negative extreme deviations |  |  |  | Positive extreme deviations |  |  |  |
| --- | --- | --- | --- | --- | --- | --- | --- | --- |
| | Participants with deviation, % | Max overlap, % | Significant voxels at $p < .05$ ($p_{FDR} < .05$), % | | Participants with deviation, % | Max overlap, % | Significant voxels at $p < .05$ ($p_{FDR} < .05$), % | |
|  |  |  | PAT | HC |  |  | PAT | HC |
| **HC<sub>test</sub>** | 21.56 | 1.49 |  |  | 60.59 | 5.2 |  |  |
| **ADHD** | 16.34 | 1.96 | 0 | 0 | 68.63 | 7.19 | 1.24 (0) | 0.22 (0) |
| **ASD** | 34.16 | 3.47 | 0.42 (0) | 0 | 71.78 | 5.94 | 4.31 (0) | 0.12 (0) |
| **BP** | 20.18 | 1.75 | 0 | 0 | 59.65 | 4.82 | 0.30 (0) | 3.00 (0) |
| **MDD** | 19.88 | 4.35 | 0.02 (0) | 0 | 50.93 | 4.97 | 0.06 (0) | 2.92 (0) |
| **OCD** | 13.17 | 3.59 | 0 | 0 | 64.07 | 4.19 | 0.14 (0) | 2.58 (0) |
| **SCZ** | 35.77 | 3.39 | 0.07 (0) | 0 | 63.97 | 5.22 | 1.52 (0) | 0.51 (0) |

### Voxel-scale WMV heterogeneity

Figures S2 and S5 quantify the spatial overlap across scales for the HCtest cohort for negative and positive extreme deviations, respectively. As there were fewer significant case-control differences in overlap for positive deviations (Figure S5-9), we focused on negative WMV deviations (i.e., areas where volume is lower than normative expectations). We quantified heterogeneity in WMV deviations as the proportion of individuals showing an extreme deviation in each voxel, estimated separately for each diagnostic group and the HCtest cohort. Across all groups, we found very little spatial overlap in the location of voxel-scale negative extreme deviations, with the maximum percentage overlap never exceeding 5% (Table 2, Figure 1f-g).

Comparison of overlap between patients and controls, revealed small, isolated areas (<1% of WMV voxels) of greater overlap in ASD, MDD, and SCZ that did not survive false discovery rate (FDR) (35) correction (pFDR < .05, two-tailed; Table 2, Figure 1h). No voxels showed significantly greater overlap in the HCtest group (Table 2). These results extend past findings (23,24), indicating that there is minimal overlap in person-specific extreme WMV deviations, highlighting a high degree of heterogeneity in voxel-wise WMV pathology across psychiatric disorders.

### Tract-scale WMV heterogeneity

We investigated whether heterogeneous extreme deviations in each clinical group converged on common axonal pathways linking specific brain regions. The maximum overlap across the 8646 connections linking 132 grey matter regions ranged from 4.19% (OCD) to 15.40% (SCZ), with 8.18% in the HCtest group (Table S3, Figure 2g-h).

Connections showing statistically significant differences in overlap scores are shown for each disorder in Figure 2i. We observed significantly greater overlap in tracts that were predominately connected to visual and amygdala regions for individuals with ASD, and connections to somatomotor areas for SCZ (*p*<.05 FDR-corrected, two-tailed), compared to controls (Table S3, Figure 2i). For ADHD, BP, MDD, and OCD, no connections survived FDR correction.

Controls showed some evidence of greater overlap in compared to individuals with OCD, but very few connections survived FDR correction (Table S3, Figure 2i).

Alternative analysis thresholds yielded comparable findings (see Methods, Table S4 and Figure S3). These results suggest that WMV deviations affect common axonal pathways in some disorders, although the overlap was only marginally higher than in the voxel-wise analysis, suggesting considerable heterogeneity at the tract-scale.

### Region-scale WMV heterogeneity

The maximum overlap observed in any given brain region was 14.87% for the held-out control group, and ranged between 8.98% (OCD) and 33.42% (SCZ) for the clinical groups (Table S5, Figure 3f-g). We observed significantly greater overlap across the cortex and subcortex in ASD and SCZ, compared to controls, with differences identified in 86% and 91% of regions, respectively (*pFDR* < .05, two-tailed, Figure 3h). ADHD and OCD showed evidence of significantly less overlap than controls, but these results did not survive FDR corrections (Figure 3h).

### Network-level WMV heterogeneity

Finally, we evaluated network-scale overlap scores between cases and controls across seven cortical networks and three subcortical areas. Overlap ranged from 10.18% somatomotor, dorsal attention, salience/ventral attention, and frontoparietal networks in OCD) to 35.25% (dorsal attention network and basal ganglia in SCZ). In controls, the maximum overlap was 19.70% in the dorsal attention network and basal ganglia (Figure 4a). Networks showing statistically significant differences in network-scale overlap are shown for each disorder in Figure 4b. ASD and SCZ had greater overlap in all networks compared to controls (pFDR < .05, two-tailed), while OCD showed less overlap in all networks except the medial temporal lobe compared to controls (pFDR < .05, two-tailed, Figure 4b). No significant differences were found for ADHD, BP, or MDD. Similar results were obtained with a 20-network parcellation (Table S7, Figure S4).

**Figure 4.**
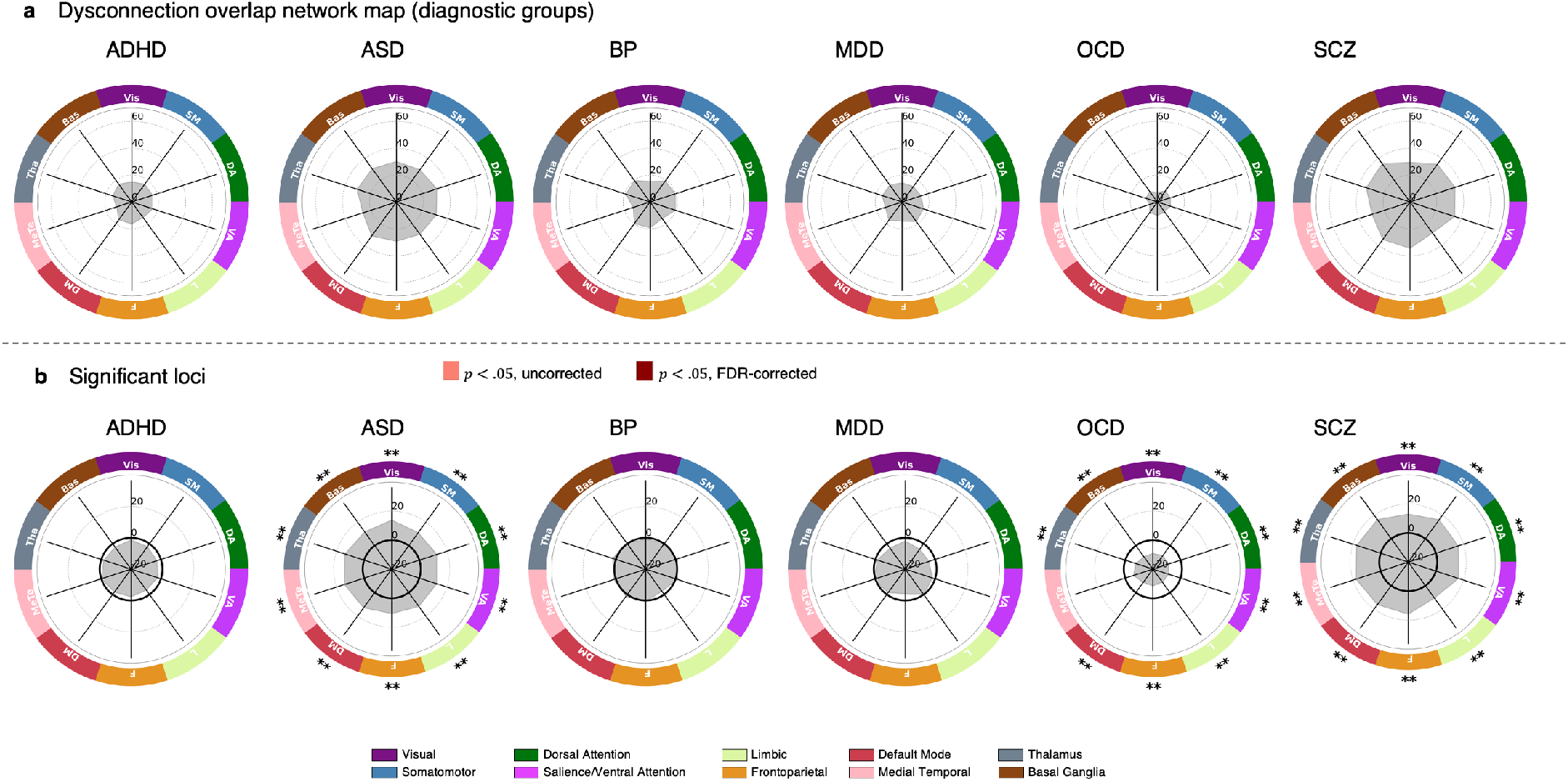
Functional networks affected by extreme negative WMV deviations. (**a**) Network maps quantifying the proportion of individuals showing at least one significant deviation in each network, for each diagnostic group. (**b**) The network-scale difference in percent overlap for extreme negative WMV deviations (z < - 2.6, cluster size threshold=10 voxels) between each clinical group and the control cohort. ** corresponds to pFDR < .05, two-tailed, * corresponds to p < .05, two-tailed. The solid black line indicates -log10 p =1.6 (p=.05, two-tailed, uncorrected). VIS, Visual; SM, Somatomotor, DA, Dorsal attention; SAL/VA, Salience/ventral attention; L, Limbic; F, Frontoparietal; DM, Default mode; MeTe, Medial Temporal; Tha, Thalamus; Bas, Basal Ganglia.

## Discussion

We characterised the structural network context of anatomically heterogeneous white matter volume (WMV) abnormalities in a diverse range of psychiatric disorders. We found that extreme heterogeneity of voxel-wise WMV abnormalities is a feature of most psychiatric disorders. For ASD and SCZ, these heterogeneous loci affected connections linked to almost all brain regions and implicating nearly all brain networks. Our findings thus suggest that WMV deviations are highly heterogeneous at the voxel and tract scales in all disorders, and only aggregate within common, yet widespread brain regions and networks, in ASD and SCZ.

At a voxel-scale, few areas showed significantly greater overlap in cases compared to controls. No voxel in any disorder showed an extreme deviation in more than 5% of cases for negative deviations and 8% of cases for positive deviations. The high degree of voxel-wise heterogeneity in WMV deviations across disorders aligns with previous normative modelling studies of WMV in SCZ, BP and ADHD (23,24) that used a subsample of the data in the current study, in addition to similar normative modelling studies of other neural phenotypes in a range of psychiatric diagnoses (19–21,25,26). The high degree of voxel-wise heterogeneity supports the contention that group-level maps of case-control differences are likely to mask considerable within-group heterogeneity.

Despite voxel-scale heterogeneity, we found that deviations aggregated within common tracts, regions, and functional networks for some disorders. This aggregation is expected as coarser scales combine data over larger regions. The HCtest sample provided a normative benchmark for expected overlap at each network scale. At the tract-scale, ASD showed greater overlap in edges linking visual areas and between visual areas and the amygdala. This aligns with ASD’s social impairments (36) and atypical visual perception (37), with research implicating amygdala dysfunction (38). Amygdala lesions have also been associated with impairments in social judgement (39), which have been likened to “acquired autism” (38,40). SCZ showed greater overlap in edges linking sensory-motor areas and connecting the sensory-motor cortex with the thalamus, consistent with previous findings of altered somatomotor-thalamic connectivity in SCZ (41–44). However, for both disorders, the maximum overlap for any connection did not exceed 15%, indicating significant individual variability.

At the region-scale, robust differences in negative deviation overlap were found for ASD and SCZ across nearly the entire brain, except for some parietal and temporal areas in the left hemisphere for ASD, and medial frontal areas and the basal ganglia for SCZ. For positive deviations in ASD, differences were observed across most of the brain, except for unimodal regions in the right hemisphere. At the network-scale, almost all functional networks were implicated in individuals with ASD or SCZ who had extreme negative deviations (34% and 36% of participants, respectively), and all individuals with ASD who had extreme positive deviations (72% of participants). These findings support evidence of widespread connectivity changes in both conditions (45–47) and align with reports of high comorbidity and overlaps in genetics, neuroimaging, clinical signs, and cognitive features between ASD and SCZ (48).

In OCD, all functional networks were less likely to be implicated compared to controls, which may simply reflect a lower total WMV deviation burden in the disorder. While past work suggests that disruption of frontostriatal systems is linked to OCD (49), our work aligns with a recent study reporting that altered structural connectivity is unlikely to drive the observed functional dysregulations (50).

We found no significant differences in WMV overlap between controls and ADHD, BP, or MDD, suggesting that WMV aberrations may be less pronounced in these conditions compared to SCZ or ASD. Indeed when taken together with our previous work examining grey matter volume (22), the current findings suggest that WMV deviations, as assessed by VBM, may be relatively sparse in psychiatric disorders. Less than 36% of individuals in any group showed at least one extreme negative WMV deviation, compared to 76% for grey matter volume deviations in our prior study (22). Positive WMV deviations were more common, with over 61% of people showing at least one extreme deviation, consistent with previous research (23,24). It remains unclear if this increase in WMV is a true biological phenomenon or a limitation of current normative models in capturing non- linearities in the voxel-wise WMV estimates. Using more flexible normative models (30,51) might help address this issue.

### Limitations and conclusions

The quantification of WMV deviations was based on our previous work on grey matter volume deviations, using the Jacobian of the non-linear warp field. However, unlike grey matter, white matter’s fibrous bundles have a strong orientation dependence. Changes perpendicular to fibre orientations indicate modifications in axon number or calibre, while changes along fibre orientations mainly affect fibre length, which has a weaker impact on information transfer. Combining registration tailored to white matter with tractography may provide greater specificity (52,53).

Diagnoses based on nosological systems such as the DSM (36) are clinical heuristics that do not necessarily define biological entities, with several well-documented limitations including pervasive comorbidity (54) and within-group clinical heterogeneity (11). As per our previous work (22), we used DSM diagnoses to evaluate heterogeneity and explore potential neural correlates of phenotypic similarities. However, this reliance may limit observed neural overlap. Focusing on syndromes that cut across diagnostic boundaries (55) might yield more precise mappings of voxel-wise deviations, their network context, and behaviour. Group differences in total deviation burden can drive differences in group overlap, complicating inferences about specific targeting of connections, regions, or networks. While grey matter volume studies have methods to address this (22), WMV analyses lack such null models. Therefore, we cannot determine if observed overlaps reflect differences in deviation burden or selective targeting, although these differences still have phenotypic consequences (see (22) for a detailed discussion).

In summary, our analysis across six psychiatric disorders found very little overlap among individuals with the same disorder in the location of MRI-based WMV deviations. For ASD and SCZ, these deviations affect common axonal pathways linking pairs of brain regions and distributed brain networks. However, the prevalence of WMV deviations is lower than prior observations for grey matter volume using the same sample (22), suggesting that WMV may be relatively spared in psychiatric illness.

## Acknowledgements

CRediT Statement Ashlea Segal: conceptualization, methodology, software, formal analysis, data curation, writing – original draft, writing – review & editing. Robert E Smith: methodology, software, writing – review & editing. Sidhant Chopra: methodology, formal analysis, writing – review & editing. Stuart Oldham: methodology, formal analysis, writing – review & editing. Linden Parkes: investigation, methodology, writing – review & editing. Kevin Aquino: methodology, writing – review & editing. Seyed Mostafa Kia: methodology, software, writing – review & editing. Thomas Wolfers: methodology, software, writing – review & editing. Barbara Franke: investigation, writing – review & editing. Martine Hoogman: investigation, writing – review & editing. Christian F. Beckmann: investigation, writing – review & editing. Lars T. Westlye: investigation, writing – review & editing. Ole A. Andreassen: investigation, writing – review & editing. Andrew Zalesky: investigation, writing – review & editing. Ben J. Harrison: investigation, writing – review & editing. Christopher G. Davey: investigation, writing – review & editing. Carles Soriano-Mas: investigation, writing – review & editing. Narcís Cardoner: investigation, writing – review & editing. Jeggan Tiego: investigation, writing – review & editing. Murat Yücel: investigation, writing – review & editing. Leah Braganza: investigation, writing – review & editing. Chao Suo: investigation, data curation, writing – review & editing. Michael Berk: investigation, writing – review & editing. Sue Cotton: investigation, writing – review & editing. Mark A. Bellgrove: investigation, writing – review & editing, funding acquisition. Andre F. Marquand: methodology, software, resources, data curation, writing – review & editing. Alex Fornito: conceptualization, methodology, formal analysis, resources, writing – original draft, writing – review & editing, supervision, funding acquisition

## Data availability

Summary of data availability for each dataset used in this study is described below. Autism Brain Imaging Data Exchange I (ABIDE I)(56) and ABIDE II(57) datasets are available through the ABIDE repository, http://fcon_1000.projects.nitrc.org/indi/abide/. The Australian Schizophrenia Research Bank (ASRB)(58) dataset is available through the ASRB repository, subject to approval of the ASRB Access Committee https://www.neura.edu.au/discovery-portal/asrb/. First Episode Mania Study (FEMS),(59) Monash Cohort (MON),(60) Obsessive-compulsive and problematic gambling study (OCDPG),(61) SPAINOCD,(62) and YoDA(63) datasets are available from the principal investigators of the respective studies, subject to evaluation of the request and local ethics committee requirements. The International Multi-centre persistent ADHD CollaboraTion (IMpACT-NL)(64) and TOP15(65) datasets are not publicly available due to privacy or ethical restrictions. Kansas Musical Depression Study (KANMDD; ds000171),(66,67) Massachusetts Institute of Technology Autism Study (MITASD; ds000212.v1.0.0),(68,69) Russia fMRI Depression Study (RUSMDD; ds002748.v1.0.5),(70,71) University of Washington ASD Study (WASHASD; ds002522.v1.0.0)(72,73) datasets are available through the OpenNeuro repository, https://doi.org/10.18112/openneuro). Human Connectome Project (HCP) dataset is available in the Human Connectome Project repository, https://www.humanconnectome.org/study/hcp-young-adult.

