## Supplementary Materials for "Multiscale heterogeneity of white matter morphometry in psychiatric disorders"

### Supplementary Methods

#### Participants

This study included 3746 individuals (1865 healthy controls, HC; 1833 cases across six different diagnostic categories) from 14 separate, independently acquired dataset and 25 scan sites. Full details on study design and clinical characteristics have been described previously for each dataset (see Table 1 for relevant references). Ethnicity was not reported in many of the datasets, and for those datasets that did report ethnicity, most, if not all participants were Caucasian. Each study was approved by the relevant ethics committee and written informed consent was obtained from each participant.

The final sample we examined was taken from a larger pool of individual recruited across the 14 datasets. In addition to the specific quality control procedures used within each study (references in Table 1), we performed a series of additional quality control checks and exclusions for our analysis. Specifically, we excluded participants who:

- Were below 18 years or above 64 years of age (*N*=346)
- Lacked the necessary clinical data (such as a clinical diagnosis or, for healthy controls, the absence of any clinical diagnosis) or demographic information (age and sex, *N*=53)
- T1-weighted structural magnetic resonance imaging (MRI) scans failed the preprocessing pipeline (*N*=63)
- Had T1-weighted MRI scans that did not pass our rigorous manual (*N*=53) and automated quality control procedures (*N*=153), as detailed in Segal et al. (1)
- Were from sites with fewer than 10 individuals in the same group and sex (described in the Normative model section below; *N*=217)

Thus, our final sample for analysis included 2,759 individuals (1,465 controls and 1,294 cases). Table 1 presents demographic details of this cohort.

#### Anatomical data

Quality assurance procedures and the data processing pipeline, which was applied to all raw T1-weighted images obtained for each dataset, are provided in detail in Segal et al. (1), with image acquisition parameters for each dataset provided in Table S1 in Segal et al. (1).

Briefly, voxel-wise WMV was estimated using the Computational Anatomy Toolbox(2) (CAT12 r113, http://dbm.neuro.uni-jena.de/cat/), which is included as an extension of Statistical Parametric Mapping software (SPM12, http://www.fil.ion.ucl.ac.uk/spm/software/spm12) in MATLAB v9.8 using the default settings.

First T1-weighted images were corrected for bias-field inhomogeneities, segmented into grey matter, white matter, and cerebrospinal fluid, then spatially normalised to the SPM MNI IXI555 template (https://brain-development.org/ixi-dataset/). The normalised segmented white matter images were modulated and spatially smoothed with an 8mm full width at half maximum (FWHM) Gaussian smoothing kernel. This approach allowed us to assess voxel-wise differences in the absolute amount (volume) of white matter, taking the extent of voxel-wise volumetric adjustments needed to align each participant with the registration template into account. Finally, to exclude non-white matter voxels from our analysis, we generated a mean image from all the normalized unmodulated white matter maps and retained voxels with a tissue probability ≥ 0.2.

#### Normative modelling

Normative models estimate the centiles of variation around the mean (referred to as the normative range) of a response variable such as WMV from a set of clinically relevant covariates, such as age and sex, across a large control sample (defined here as a group of people without a psychiatric diagnosis), referred to as a *training set*. These estimates are then used to quantify deviations from the normative range of individuals in the *test set,* which typically consists of cases of interest and a held-out set of controls for comparison.(3–5)

As per Segal et al.(1), the training set ($HC_{train}$, *n* = 1,465) for the normative model was created by randomly selecting from each scanner site either 90% of controls or all controls if the sample size for that site was less than 30. The test set consisted of all clinical data (*n* = 1,294), as well as the remaining controls from each scan site ($HC_{test}$, *n* = 269), which offered a normative benchmark for assessing case-specific model deviations.

B-spline basis expansion over age was used to model non-linear effects, and likelihood warping was used to model non-Gaussian and heteroscedastic imaging phenotypes. To handle the site effect, we included scan site as a covariate (6), as in (7).

To identify positive and negative extreme deviations in white matter voxel (WMV) estimates from the normative model, we thresholded the deviation maps at $|z| > 2.6$ ($p < .005$) and applied a 10-voxel extent threshold (Figure 1d). This method was used for each individual in both clinical and control ($HC_{test}$) groups. We chose this approach for consistency with previous work (1,8) and because it provides a uniform definition of extreme deviations, unlike adaptive thresholding methods like the false discovery rate (FDR) (9), which vary per individual.

#### Evaluation of model performance

We assessed model fit for each brain region by evaluating five performance metrics: (i) explained variance (EV); (ii) the standardized mean-squared error (SMSE); (iii) the mean standardized log-loss (MSLL); (iv) skew; and (v) kurtosis. We removed voxels with poor performance (SMSE > 1.5, MSLL > 0.5, or skew > 1.5), which amounted to 1.7% voxels in total (Figure S1). We also evaluated the model's efficacy in partitioning site-related variance in the data using linear support vector machines. Specifically, we used a series of one-versus-all linear support vector machines (with default slack parameter = 1) trained on the deviation z-maps from the $HC_{test}$ subset to classify scan sites. For each site, we ran a 2-fold SVM classifier to obtain the mean balanced accuracy score for the given site. In this analysis, a balanced accuracy close to chance-level (50%) indicates that the resulting deviations were not contaminated by residual site effects (Table S2)

#### Characterizing voxel-wise heterogeneity of extreme deviations

We characterized the heterogeneity of thresholded person-specific extreme deviation maps for each disorder using a non-parametric approach, following Segal et al. (1). This procedure involved subtracting the $HC_{test}$ overlap map from each disorder's overlap map, resulting in an overlap difference map for each disorder, separately for positive and negative extreme deviations. For each disorder, we then permuted group labels (i.e., HC, case) and repeated the procedure 10,000 times to derive an empirical distribution of overlap difference map under the null hypothesis of random group assignment. For each white matter voxel, we obtained $p$-values as the proportion of null values that exceeded the observed difference. The tails of the null distribution (i.e., values associated with $p<0.10$) were approximated using a generalized Pareto distribution (10), as implemented in the Permutation Analysis of Linear Models software package (PALM alpha116) (11), to allow inference at arbitrarily high levels of precision. Statistically significant effects were identified using an FDR-corrected (9) threshold of $p_{FDR}<.05$, two-tailed.

#### Diffusion-weighted imaging acquisition parameters and processing

We used diffusion tractography to identify the axonal pathways that intersected each WMV deviation (Figure 1f-h). We then used this information to build matrices encoding the specific tracts affected by WMV deviations (Figure 1i-k). The deviation-related structural circuitry was mapped in an independent cohort as high-quality DWI data is essential for accurate estimate of tractograms and such data was not available for all clinical samples. This logic precisely aligns with the logic of standard lesion network mapping studies of neurological patients (12–16). We focused on the same sub-sample of 150 people in the HCP dataset as Segal et al (1) to minimise the computational burden.

Data were acquired on a customized Siemens 3 T Connectome Skyra scanner at Washington University in St. Louis, MO, USA, using a multishell protocol for the diffusion weighted imaging with the following parameters: 1.25-$mm^3$ voxel size; repetition time (TR) = 5520 ms; echo time (TE) = 89.5 ms; field of view (FOV) of 210 mm by 180 mm; 270 directions with $b$ = 1000, 2000, 3000 s/$mm^2$ (90 per $b$ value); and 18 $b$ = 03 volumes. Structural T1-weighted data were acquired with 0.7-mm voxels, TR = 2400 ms, TE = 2.14 ms, and an FOV of 224 mm by 224 mm.

The diffusion data used underwent the HCP minimal pre-processing pipeline,(17) which included normalization of mean $b$ = 0 images across diffusion acquisitions and correction for echo-planar imaging susceptibility and signal outliers, eddy current–induced distortions, slice dropouts, gradient nonlinearities, and participant motion. T1-weighted data were corrected for gradient and readout distortions before being processed with FreeSurfer.(18) The details of this pipeline are provided in more detail elsewhere.(19) Using the corrected diffusion data, we estimated fibre orientation distributions (FODs) using multishell Constrained Spherical Deconvolution,(20) which formed the basis for probabilistic tractography with the Fibre Orientation Distributions (iFOD2) algorithm, as implemented in MRtrix3.(21–24) We applied Anatomically Constrained Tractography (ACT) to improve the biological accuracy of the structural networks.(25) ACT uses multi-tissue segmentation to ensure streamlines are beginning, traversing, and terminating in anatomically plausible locations. A total of 10 million streamlines were generated using dynamic seeding along with default MRtrix3 parameters.(21–24,26)

#### Streamline region assignments

We examined streamline estimates of connectivity between 132 regions, defined by combining well-validated parcellations of the cortex (Schaefer 100 region 7 network parcellation)(27) and subcortex (Tian Subcortical Atlas Scale II, 32 region parcellation).(28) The atlas was then transformed from the template surface to the surface of each individual in the $HCP_{150}$ sample based on the spherical registration procedure implemented in FreeSurfer.(18) Next, the parcellation was projected to a volumetric image and resampled to the same resolution as the $HCP_{150}$ diffusion data using *FSL's applywarp* with nearest-neighbour interpolation.(29) The parcellation was subsequently combined with each person's tractogram and streamlines were assigned to the nearest brain region within 5 mm of the streamline endpoints using *MRtrix3* (v 3.0) *tck2connectome* function.(24,30)

The tractograms were transformed to MNI IXI555 space to facilitate interpretation with respect to the individual-specific WMV deviation maps. To this end, we used *FSL(v 6.0.6) FLIRT* and *FNIRT*(29,31) to generate and compose transforms for co-registering each $HCP_{150}$ participant's diffusion data to their T1-weighted volume, and their T1-weighted volume to the SPM MNI IXI555 template. Finally, we used *MRtrix's tcktransform* to warp the tractogram to MNI IXI555 space using the composed warp.

#### Identifying streamlines intersecting with WMV deviations

To construct the dysconnection overlap matrices, we used the extreme WMV deviation cluster map (Figure 2a) of each patient/control to filter each $HCP_{150}$ individual's tractogram (Figure 2b-c) to retain only streamlines intersecting with at least one WMV cluster from the deviation maps. This generated a filtered streamline count matrix ($C'$), indicating pairwise connections affected by deviations. We binarized $C'$, setting $C'_{ij}$ to 1 if the filtered edge's streamline count exceeded 10, and 0 otherwise (Figure 2d).

The $C'$ matrices were aggregated across $HCP_{150}$ individuals to create a consensus matrix ($C^G$), which encoded the proportion of participants for whom each edge was present in $C'$. This matrix estimates the probability that a given connection is associated with observed WMV deviations in that participant in the clinical group (or $HC_{test}$ group), given individual variability in normative connectome architecture. A separate $C^G$ matrix was obtained for each individual in the clinical and $HC_{test}$ groups. We binarized $C^G$, setting edges with weights less than 50% to zero, to encode connections affected by a WMV extreme deviation (termed extreme edge deviation matrix; Figure 2e). We repeated the analysis with a 75% threshold to validate our findings (see Figure S3).

Finally, we aggregated these binary $C^G$ matrices for each clinical and $HC_{test}$ group, separately for positive and negative deviations, creating group-specific overlap matrices. Each element in these matrices shows the proportion of individuals in each group affected by an extreme deviation in that pairwise connection, referred to as the dysconnection overlap matrix (Figure 2f).

To evaluate group differences using permutation-based inference, similar to the voxel-scale deviation analysis, we subtracted the $HC_{test}$ group's dysconnection overlap matrix from each clinical group's matrix and performed mass univariate testing across connectome edges.

Significant differences in dysconnection overlap were identified using an empirical null distribution from 10,000 permutations and a generalized Pareto tail approximation (10), with a threshold of pFDR < .05, two-tailed. This was done separately for positive and negative extreme deviations.

### Supplementary Results

#### Model evaluation

The results from the linear support vector machine indicated that the balanced accuracy scores ranged between 49.71 and 58.53% across all sites, suggesting that resulting deviations were not contaminated by residual site effects (Table S2).

Rank correlations indicated that scan quality (as measured by CAT12's IQR) was weakly associated with extreme deviation burden for the cases (*rho* = 0.07, *p*=.02), but not for $HC_{test}$ (*rho* = -0.04, *p*=.55), however this difference was not statistically significant (Fisher's z-transformation, $z = 1.64$, $p = 0.10$).

#### Analysis of positive deviations

Despite both cases and controls generally showing more extreme positive than negative deviations (Figure S6), there were few differences in the extent of overlap, with <1% of edges show significantly greater overlap in ASD and SCZ (Figure S7) compared to controls, and 76.52% of regions and all networks showing significantly greater overlap in ASD, relative to controls (Figure S8 and Figure 9 respectively). No other disorders at any of the spatial scales showed significantly greater overlap compared to controls after correcting for multiple comparisons.

### Supplementary Tables

**Table S2. Balanced classification accuracy scores (%) from the linear support vector machines from the held-out controls and cases**

| Dataset | Site | Held-out controls ($HC_{test}$) |
|---|---|---|
| ABIDE I | CALTECH | N/A |
| | CMU | N/A |
| | LEUVEN_1 | N/A |
| | MAX-MUN | 50.00 |
| | NYU | 50.00 |
| | PITT | N/A |
| | SBL | N/A |
| | USM | 50.00 |
| ABIDE II | BNI | 50.00 |

| | | |
|---|---|---|
| | IU | N/A |
| ASRB | BRIS | 50.00 |
| | MELB | 54.81 |
| | PERT | N/A |
| | SYDN | 50.00 |
| FEMS | | 50.00 |
| MON | | 56.42 |
| IMPACT | | 58.53 |
| KANMDD | | N/A |
| MITASD | | 50.00 |
| OCDPG | | 50.00 |
| RUSMDD | | 50.00 |
| SPAINOCD | | 49.71 |
| TOP15 | | 51.44 |
| WASHASD | | 50.00 |
| YoDA | | 56.25 |

*N/A = collection sites where data for HCs was < 30, therefore all HC data was included in training set.

**Table S3. Deviation burden at the tract-scale**

| | Negative extreme deviations | | | | Positive extreme deviations | | | |
|---|---|---|---|---|---|---|---|---|
| | Participants with deviation, % | Max overlap, % | Significant loci at *p<.05* ($p_{FDR}$<.05), % | | Participants with deviation, % | Max overlap, % | Significant loci at *p<.05* ($p_{FDR}$<.05), % | |
| | | | PAT | HC | | | PAT | HC |
| **HC$_{test}$** | 19.7 | 8.18 | | | 58.36 | 30.86 | | |
| **ADHD** | 16.34 | 6.54 | 0 | 0.31 (0) | 67.97 | 35.95 | 0.32 (0) | 0.16 (0) |
| **ASD** | 31.68 | 11.88 | 37.89 (6.08) | 0 | 70.79 | 40.1 | 17.26 (0.02) | 0 |
| **BP** | 19.74 | 6.14 | 0.60 (0) | 0.01 (0) | 58.77 | 32.02 | 0.18 (0) | 1.83 (0) |
| **MDD** | 19.25 | 6.21 | 0.21 (0) | 0 | 50.93 | 22.36 | 0 | 7.72 (0.01) |
| **OCD** | 10.18 | 4.19 | 0 | 1.86 (0) | 62.87 | 29.94 | 0 | 4.18 (0) |
| **SCZ** | 35.77 | 15.4 | 19.99 (3.33) | 0 | 63.19 | 39.16 | 2.59 (0.10) | 0 |

**Table S4. Deviation burden at the tract-scale (weight threshold 75%)**

| | Negative extreme deviations | | | | Positive extreme deviations | | | |
|---|---|---|---|---|---|---|---|---|
| | Participants with deviation, % | Max overlap, % | Significant connections at *p<.05* ($p_{FDR}$<.05), % | | Participants with deviation, % | Max overlap, % | Significant connections at *p<.05* ($p_{FDR}$<.05), % | |
| | | | PAT | HC | | | PAT | HC |
| **HC$_{test}$** | 19.7 | 6.69 | | | 58.36 | 28.62 | | |
| **ADHD** | 16.34 | 6.54 | 0.01 | 0.14 (0) | 67.97 | 33.99 | 0.17 (0) | 0.21 (0) |
| **ASD** | 31.68 | 10.40 | 27.68 (1.21) | 0 | 70.3 | 37.13 | 11.41(0) | 0 |
| **BP** | 19.74 | 4.82 | 0.71 (0) | 0 | 58.77 | 29.39 | 0.13 (0) | 1.62 (0) |
| **MDD** | 19.25 | 5.59 | 0.26 (0) | 0 | 50.93 | 19.88 | 0 | 5.60 (0.03) |
| **OCD** | 10.18 | 3.59 | 0 | 0.69 (0) | 62.87 | 25.15 | 0 | 3.83 (0) |
| **SCZ** | 35.77 | 13.84 | 12.53 (0) | 0 | 63.19 | 36.55 | 1.65 (0.06) | 0.05 (0) |

**Table S5. Deviation burden at the region-scale**

| | Negative extreme deviations | | | | Positive extreme deviations | | | |
|---|---|---|---|---|---|---|---|---|
| | Participants with deviation, % | Max overlap, % | Significant regions at *p<.05* ($p_{FDR}$<.05), % | | Participants with deviation, % | Max overlap, % | Significant regions at *p<.05* ($p_{FDR}$<.05), % | |
| | | | PAT | HC | | | PAT | HC |
| **HC$_{test}$** | 19.7 | 14.87 | | | 58.36 | 53.53 | | |
| **ADHD** | 16.34 | 15.03 | 0 | 1.51 (0) | 67.97 | 60.13 | 10.61 (0 | 0 |
| **ASD** | 31.68 | 27.23 | 86.36 (84.85) | 0 | 70.79 | 64.85 | 83.33 (75) | 0 |
| **BP** | 19.74 | 17.11 | 0 | 0 | 58.77 | 56.58 | 0 | 0 |
| **MDD** | 19.25 | 13.04 | 0 | 0 | 50.93 | 46.58 | 0 | 30.30 (0) |

| | | | | | | | | |
|---|---|---|---|---|---|---|---|---|
| **OCD** | 10.18 | 8.98 | 0 | 31.82<br>1.51 | 62.87 | 58.08 | 0 | 0 |
| **SCZ** | 35.77 | 33.42 | 92.42(90.91) | 0 | 63.19 | 59.79 | 8.33 (0) | 0 |

**Table S6. Deviation burden at the functional network-scale**

| | **Negative extreme deviations** | | | | **Positive extreme deviations** | | | |
|---|---|---|---|---|---|---|---|---|
| | Participants with deviation, % | Max overlap, % | Significant networks at *p<.05* ($p_{FDR}$<.05), % | | Participants with deviation, % | Max overlap, % | Significant networks at *p<.05* ($p_{FDR}$<.05), % | |
| | | | PAT | HC | | | PAT | HC |
| **HC$_{test}$** | 19.7 | 18.86 | | | 58.36 | 57.25 | | |
| **ADHD** | 16.34 | 16.34 | 0 | 0 | 67.97 | 66.67 | 0 | 0 |
| **ASD** | 31.68 | 31.68 | 100 (100) | 0 | 70.79 | 69.31 | 100 (100) | 0 |
| **BP** | 19.74 | 19.74 | 0 | 0 | 58.77 | 58.77 | 0 | 0 |
| **MDD** | 19.25 | 18.01 | 0 | 0 | 50.93 | 50.93 | 0 | 0 |
| **OCD** | 10.18 | 10.18 | 0 | 90 (90) | 62.87 | 61.68 | 0 | 0 |
| **SCZ** | 35.77 | 35.25 | 100 (100) | 0 | 63.19 | 63.19 | 0 | 0 |

**Table S7. Deviation burden at the functional network-scale (20 network)**

| | **Negative extreme deviations** | | | | **Positive extreme deviations** | | | |
|---|---|---|---|---|---|---|---|---|
| | Participants with deviation, % | Max overlap, % | Significant networks at *p<.05* ($p_{FDR}$<.05), % | | Participants with deviation, % | Max overlap, % | Significant networks at *p<.05* ($p_{FDR}$<.05), % | |
| | | | PAT | HC | | | PAT | HC |
| **HC$_{test}$** | 19.7 | 18.96 | | | 58.36 | 57.25 | | |
| **ADHD** | 16.34 | 16.34 | 0 | 0 | 67.97 | 66.67 | 0 | 0 |
| **ASD** | 31.68 | 31.68 | 100 (100) | 0 | 70.79 | 69.31 | 95 (95) | 0 |
| **BP** | 19.74 | 19.74 | 0 | 0 | 58.77 | 58.33 | 0 | 0 |
| **MDD** | 19.25 | 16.15 | 0 | 0 | 50.93 | 50.93 | 0 | 10 (0) |
| **OCD** | 10.18 | 10.18 | 0 | 80 (80) | 62.87 | 61.68 | 0 | 0 |
| **SCZ** | 35.77 | 35.25 | 100 (100) | 0 | 63.19 | 63.19 | 0 | 0 |

### Supplementary Figures

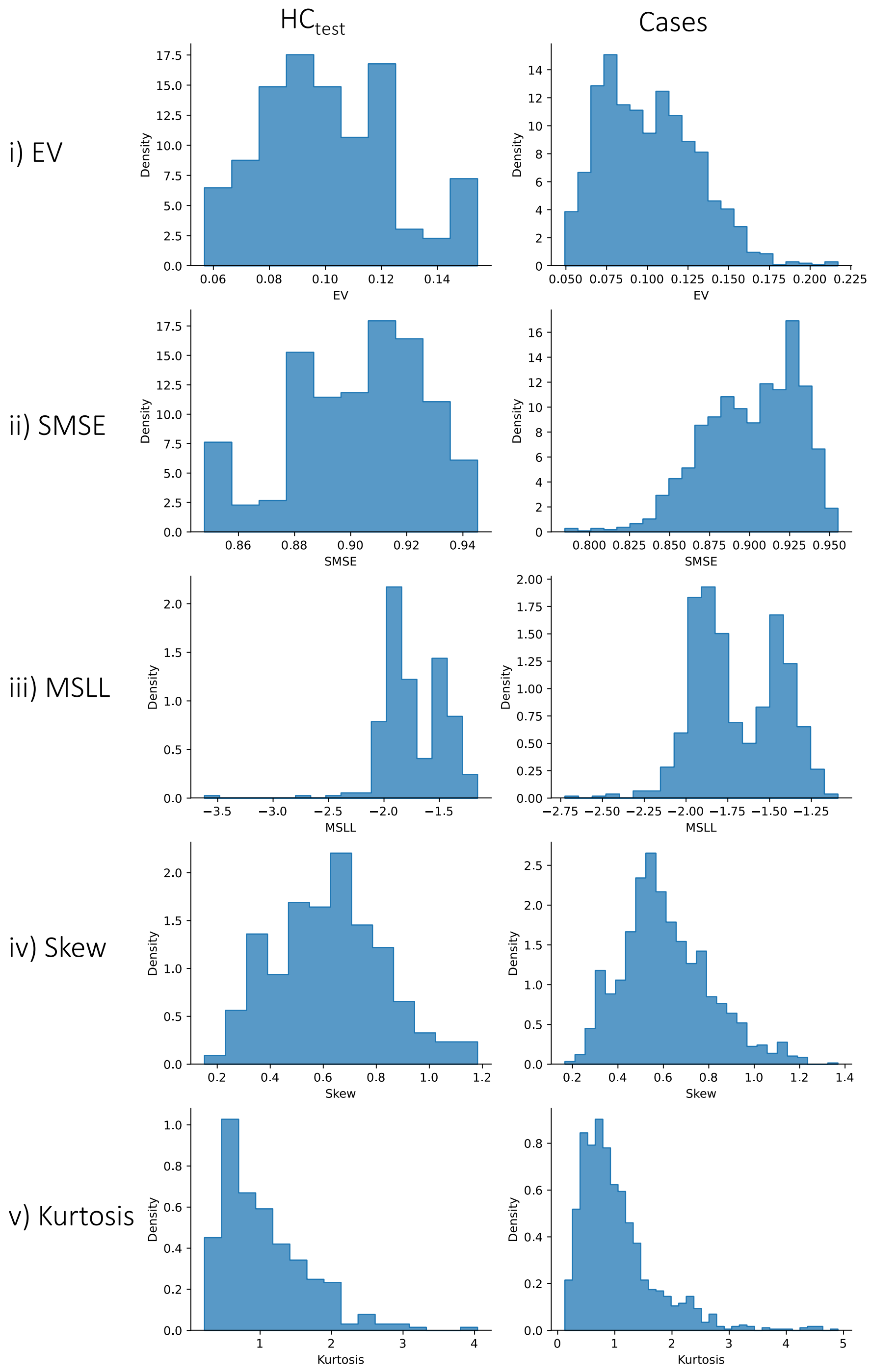

**Figure S1. Performance metrics from the normative models.** These metrics measure the accuracy with which our normative model estimated the relationship between GMV and age, sex, and site. The distributions of i) explained variance (EV; higher is better), ii) Standardized mean squared error (SMSE; lower is better), iii) Mean standardized log-loss (MSLL; lower is better), iv) Skew (close to zero is better), v) Kurtosis (close to zero is better) across WMV voxels. The left panel presents data for the HC$_{test}$ cohort, the right panel presents data for the cases.

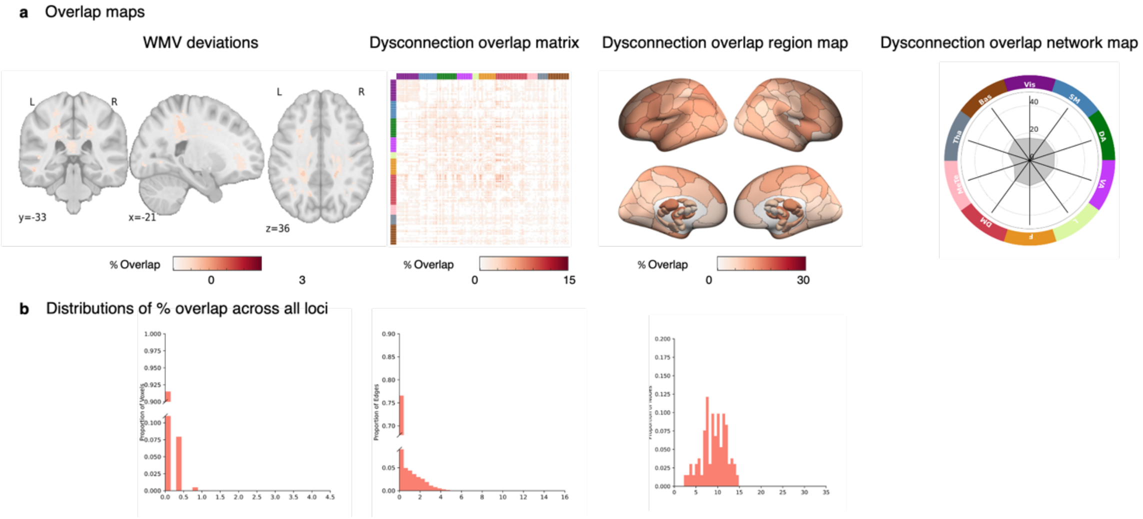

**Figure S2. Distribution of negative extreme WMV deviation overlap in $HC_{test}$ cohort across scales.**

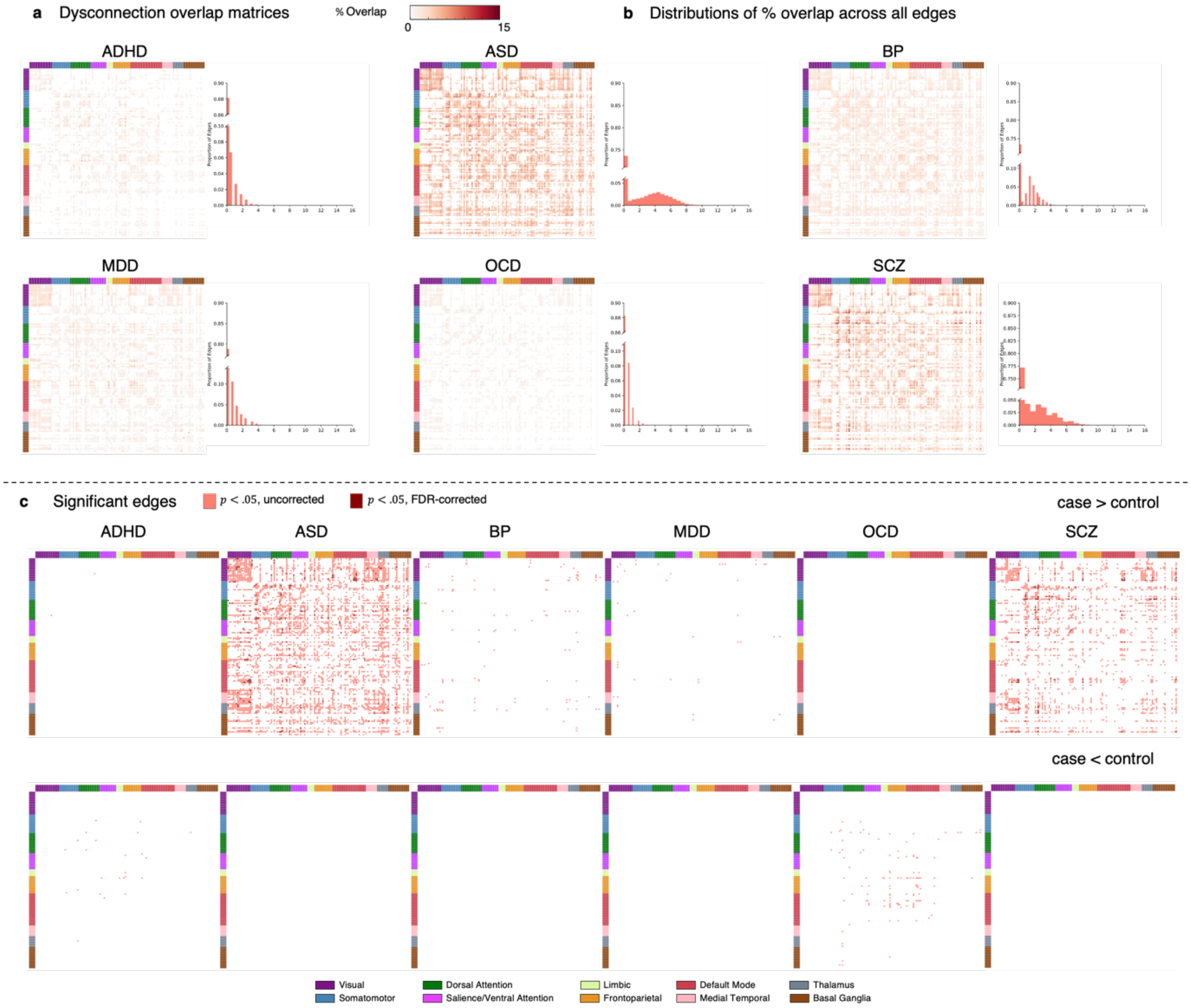

**Figure S3. Connections affected by extreme negative WMV deviations, using a weight threshold of 75%.** (a) Matrices showing the tract-scale dysconnection for each clinical group and the held-out control group ($HC_{test}$). (b) Histograms showing the distribution of overlap percentages observed across all inter-regional connections. (c) Matrices showing edges structurally connected to extreme negative WMV deviations (Z < - 2.6, cluster threshold=10) with significantly greater overlap in cases, compared to controls (top), and significantly greater overlap in controls, compared to cases (bottom, $p < .05$, two-tailed).

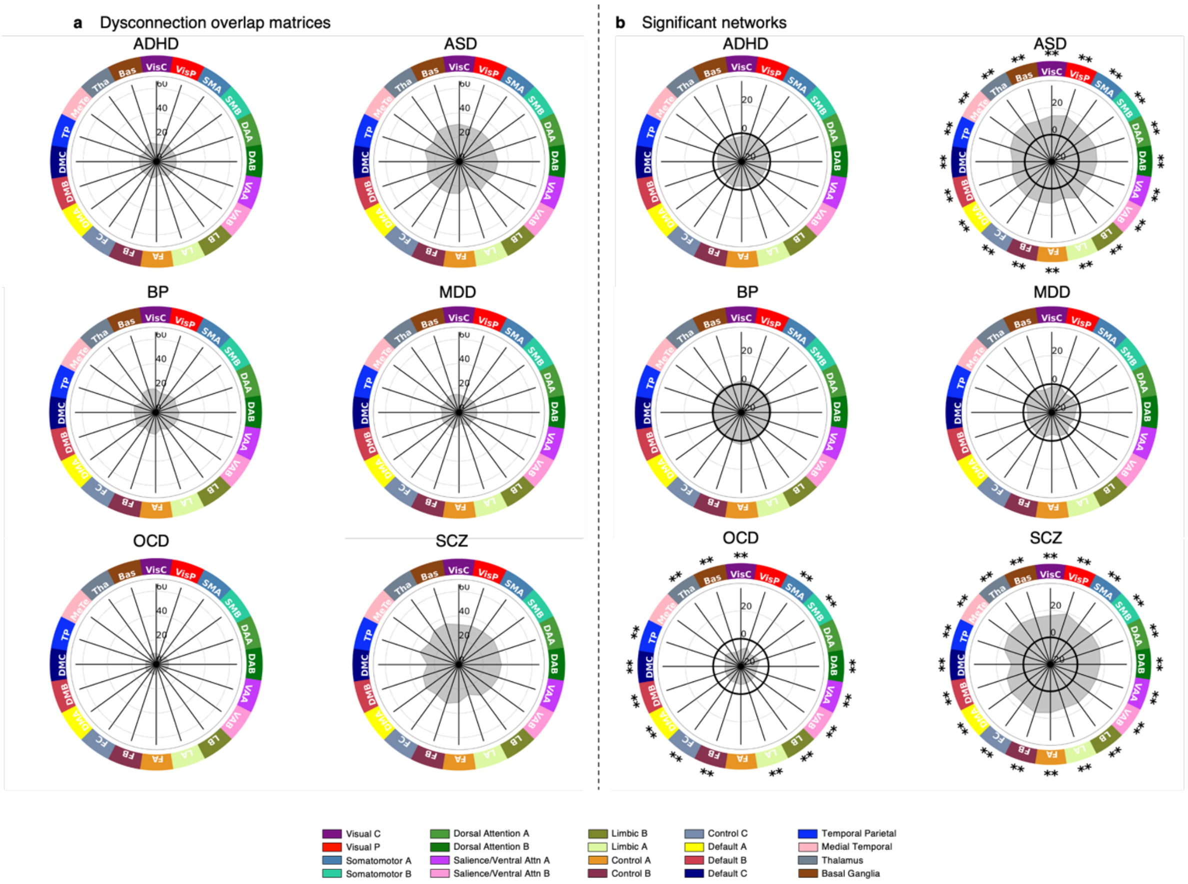

**Figure S4. Functional networks affected by extreme negative WMV deviations using 20 network parcellation.** (a) Network maps quantifying the proportion of individuals showing a significant deviation in each network, for each diagnostic group. (b) The network-scale difference in percent overlap for extreme negative WMV deviations ($z < -2.6$, cluster threshold=10) between each clinical group and the control cohort. ** corresponds to $p_{FDR} < .05$, two-tailed, * corresponds to $p < .05$, two-tailed. The solid black line indicates $-\log_{10} p = 1.6$ ($p = .05$, two-tailed, uncorrected).

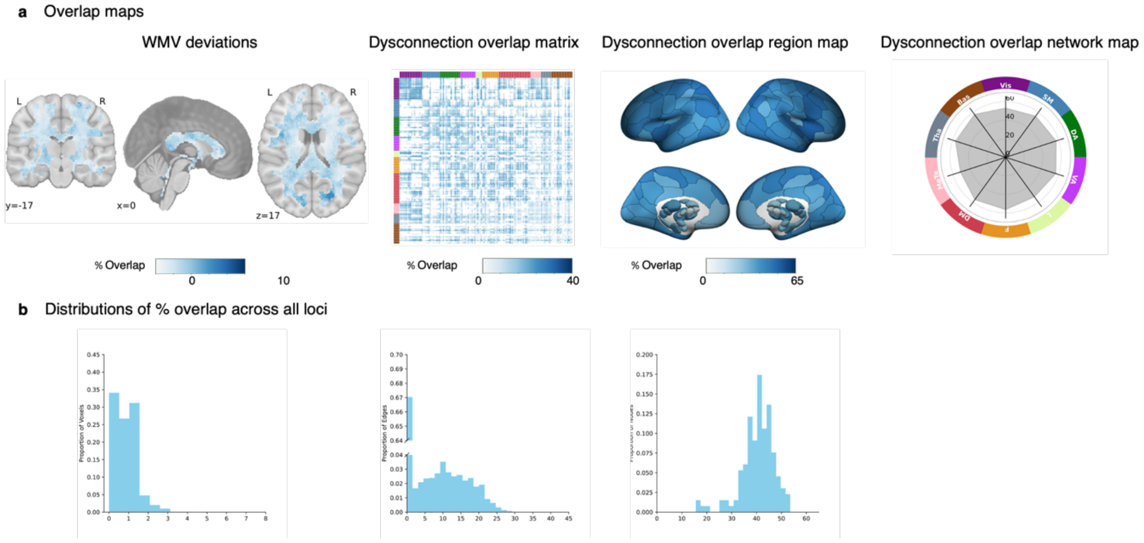

**Figure S5. Distribution of positive extreme WMV deviation overlap in $HC_{test}$ cohort across scales**

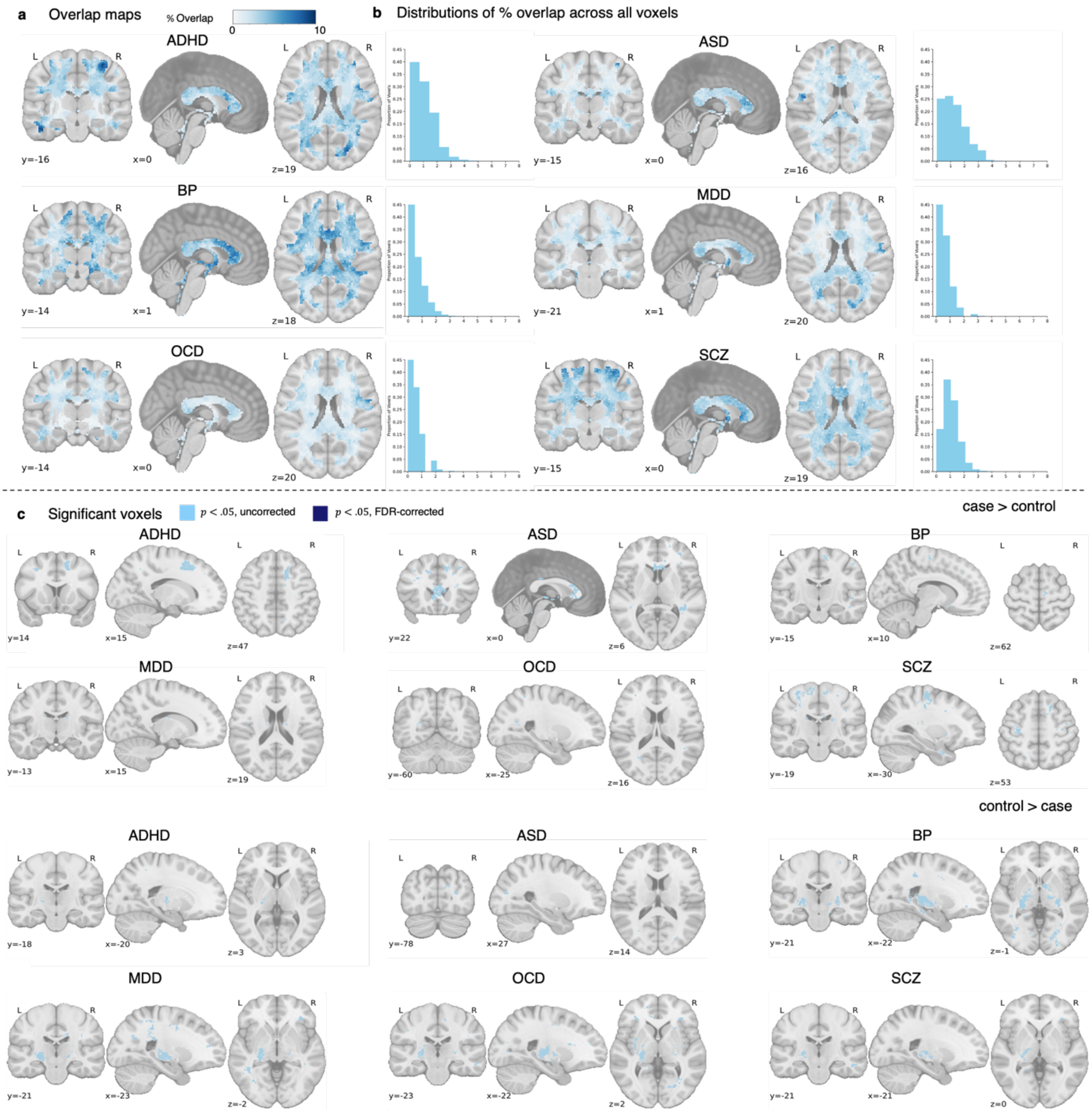

**Figure S6. Spatial overlap in extreme positive WMV deviations.** (a) Spatial map which quantifies the proportion of individuals showing significant an extreme deviation in a given voxel, yielding an extreme deviation overlap map. (b) Statistical plots showing voxels with significantly greater overlap in cases, compared to controls (left) and significantly greater overlap in controls, compared to cases in extreme positive deviations (top), and greater overlap in controls compared to cases (bottom, $p < .05$, two-tailed). $p < .05$, two-tailed)

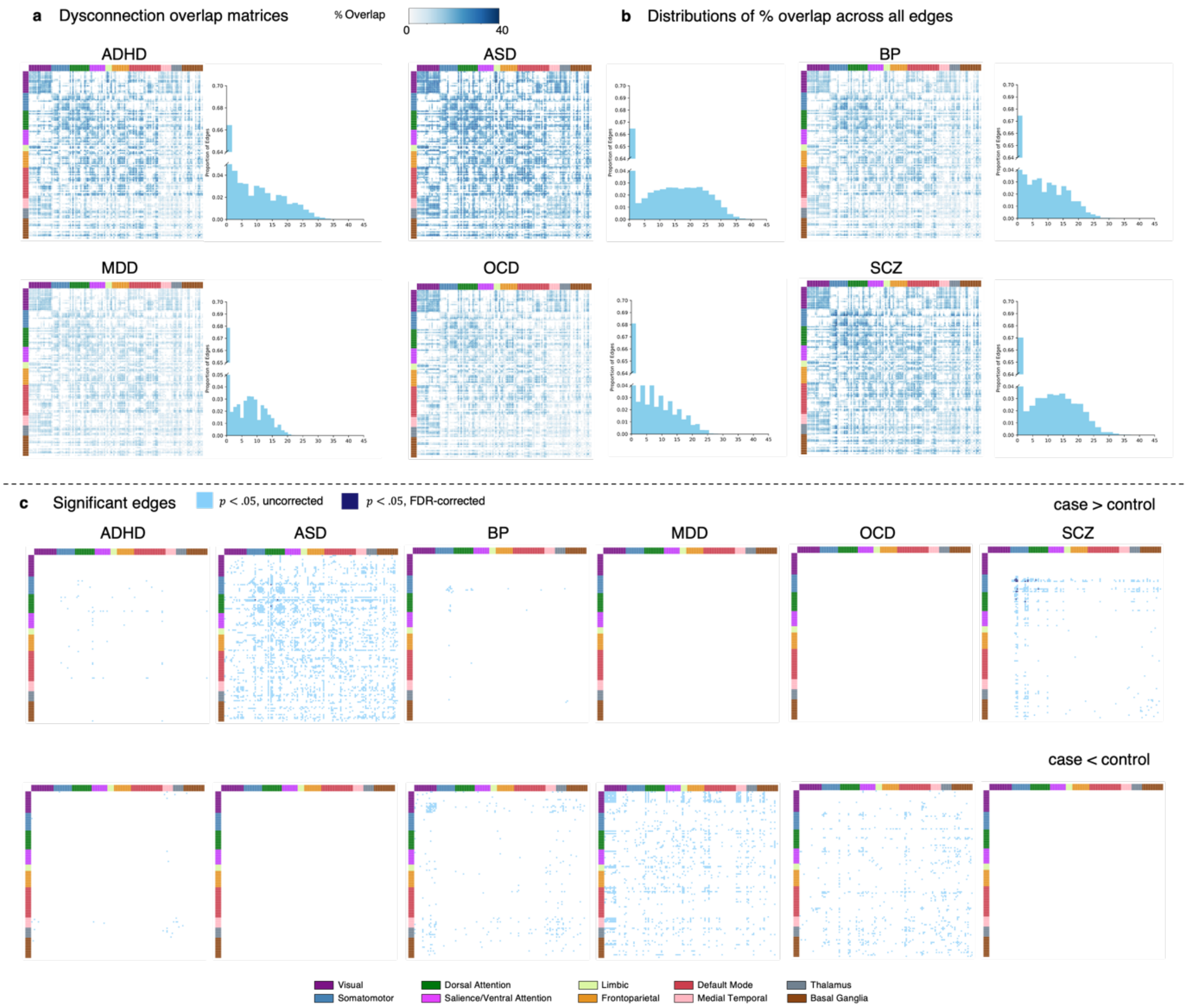

**Figure S7. Connections affected by extreme positive WMV deviations.** (a) Matrices showing the tract-scale dysconnection for each clinical group. (b) Histograms showing the distribution of overlap percentages observed across all inter-regional connections. Note that the y-axis is broken so can see the distribution of non-zero values. (c) Matrices showing edges structurally connected to extreme positive WMV deviations (Z < 2.6, cluster threshold=10) with significantly greater overlap in cases, compared to controls (top), and significantly greater overlap in controls, compared to cases (bottom, $p < .05$, two-tailed).

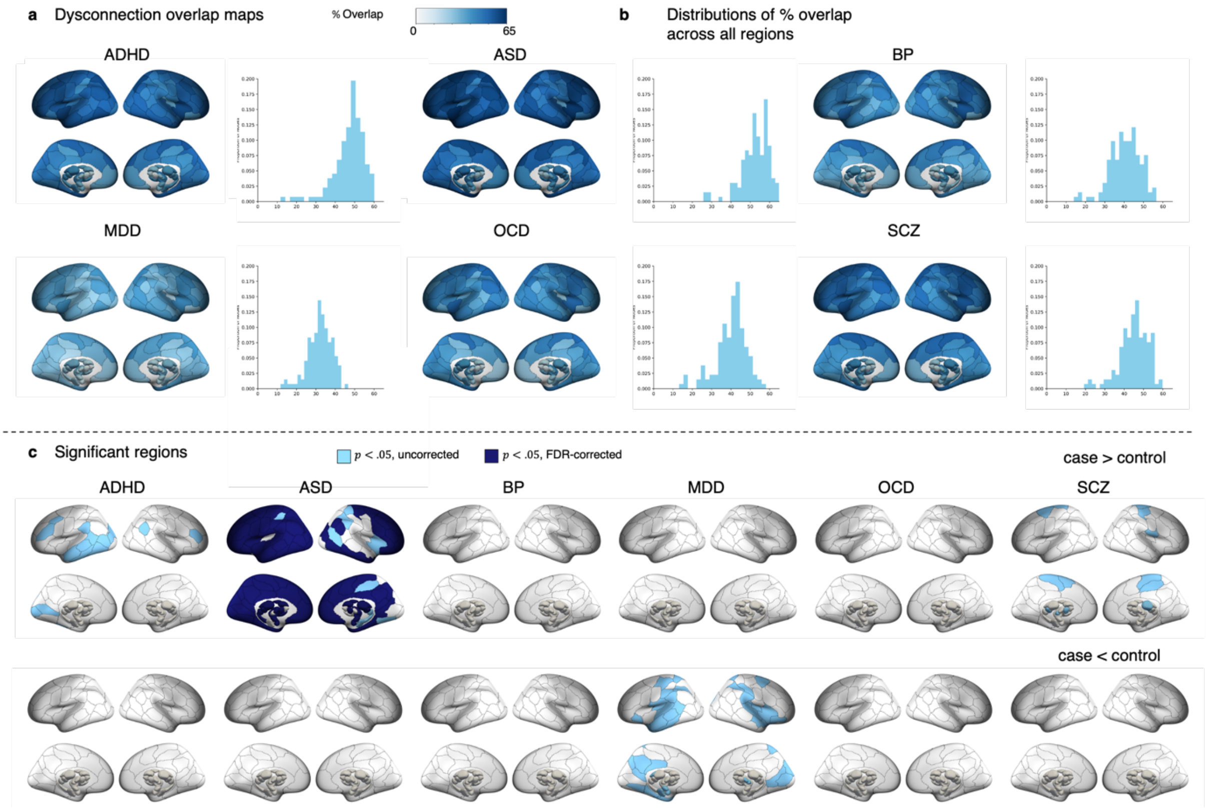

**Figure S8. Regions attached to connections affected by extreme positive WMV deviations.** (a) Spatial maps quantifying the proportion of individuals showing significant structural connectivity in each region, yielding a region-scale dysconnection map for each diagnostic group. (b) Histograms showing the distribution of overlap percentages observed across all regions. (c) Statistical maps showing regions structurally connected to extreme positive WMV deviations (z < - 2.6, cluster threshold=10) with significantly greater overlap in cases, compared to controls (top), and significantly greater overlap in controls compared to cases (bottom, $p < .05$, two-tailed).

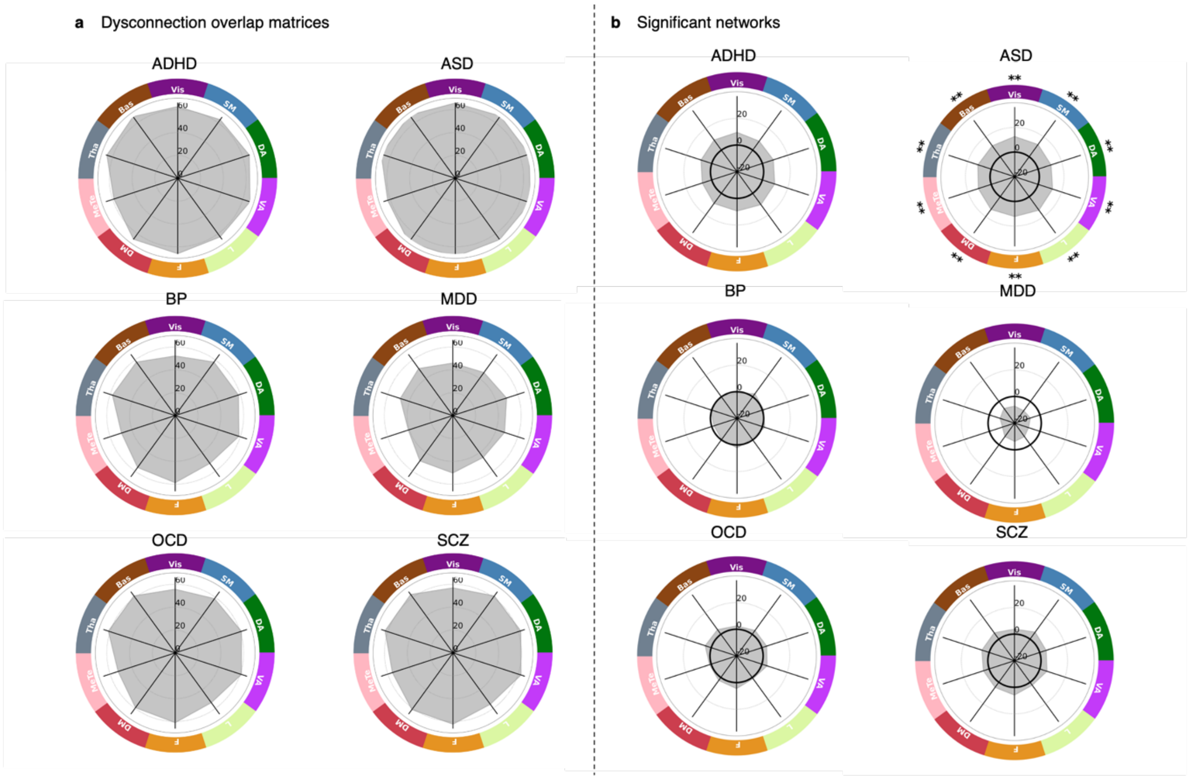

**Figure S9. Functional network affected by extreme positive WMV deviations.** (a) Network maps which quantify the proportion of individuals showing a significant deviation in each network, yielding a network-scale dysconnection map for each diagnostic group. (b) The network-scale difference in percent overlap for extreme positive WMV deviations (Z > 2.6, cluster threshold=10) between each clinical group and the control cohort. ** corresponds to $p_{FDR} < .05$, two-tailed, * corresponds to $p < .05$, two-tailed
